# Gene prediction and evolutionary characterization of heat shock transcription factors in *Salvia miltiorrhiza*

**DOI:** 10.1101/2023.05.19.541281

**Authors:** Xianwen Meng, Xiang Yan

**Affiliations:** School of Information Engineering, Yulin University, Yulin, Shaanxi, 719000, China; Shaanxi Provincial Engineering and Technology Research Center of Cashmere Goats, Yulin University, Yulin, Shaanxi, 719000, China; State Key Laboratory of Soil Erosion and Dryland Farming on the Loess Plateau, Northwest A&F University, Yangling, Shaanxi, 712100, China; State Key Laboratory of Soil Erosion and Dryland Farming on the Loess Plateau, Institute of Soil and Water Conservation, Chinese Academy of Sciences and Ministry of Water Resource, Yangling, Shaanxi, 712100, China

**Keywords:** Salvia miltiorrhiza, heat shock transcription factor genes, prediction

## Abstract

**Background:** Heat shock transcription factors (HSFs) serve as central regulators in plants and enable adaptations to dynamic environmental changes, such as heat stress. *Salvia miltiorrhiza* is an important medicinal plant model. However, various biotic and abiotic stresses, pathogen infections, and degeneration of elite cultivars limit the yield and quality of S. miltiorrhiza. Therefore, it is important to predict and characterize the stress resistance genes of *S. miltiorrhiza* and clarify potential molecular mechanisms for genetic improvement. However, the knowledge of HSF genes in *S. miltiorrhiza* is limited.

**Methods:** Thus, we performed a genome-wide analysis of HSF genes in *S. miltiorrhiza*.

**Results:** Using high-quality genome sequences, we predicted 34 HSF genes distributed unevenly across eight chromosomes. Twenty-three genes were segmentally duplicated, suggesting that in *S. miltiorrhiza*, segmental duplications have significantly contributed to the expansion of the HSF gene family. The predicted genes were divided into 15 phylogenetic groups, among which the sequence characteristics were relatively well conserved. The Ka/Ks ratios indicated that duplicated HSF genes underwent negative selection. Most of the HSF genes displayed different expression characteristics under drought stress and salicylic acid induction. This study provides valuable data that may inform further functional studies of HSF genes in *S. miltiorrhiza*.

## Introduction

*Salvia miltiorrhiza* is an important medicinal plant model that belongs to the *Salvia* genus of the Lamiaceae family (Song et al. 2013). Because of its substantial medicinal and economic value, *S. miltiorrhiza* is widely distributed and cultivated in China. Its dried roots and rhizomes, called “Danshen”, are well known in traditional Chinese medicine. *S. miltiorrhiza* is commonly used to treat cardiovascular and cerebrovascular diseases (Guo et al. 2014; Wang et al. 2017; Zhou et al. 2005). The two primary pharmaceutically active components are water-soluble phenolic acids and lipid-soluble tanshinones. These components exhibit antioxidant, antitumor, antibacterial, antiviral, and anti-inflammatory properties (Song et al. 2020; Xu et al. 2016; Zhang et al. 2015). Phenolic acids and tanshinones are secondary metabolites that are biosynthesized and accumulated in large quantities in defense against unfavorable environmental stresses. This response serves as an important evolutionary mechanism (Erb & Kliebenstein 2020). Wild resources of *S. miltiorrhiza* are extremely scarce, and the quality of cultivated resources is deteriorating; the supply of resources is failing to meet the rapidly growing market demand. Therefore, improving the quality of cultivated resources is necessary. Abiotic stresses seriously affect yield and quality. Therefore, it is important to improve the stress resistance of *S. miltiorrhiza*.

Heat shock transcription factors (HSFs) are central regulators that enable plants to respond to various biotic and abiotic stresses, including heat stress (Andrasi et al. 2021; Guo et al. 2016). HSFs mediate the expression and rapid accumulation of heat shock proteins (HSPs) by recognizing and binding to heat stress elements present in the promoters of HSP genes. Plant HSF proteins contain a typical conserved modular structure, which includes an N-terminal DNA-binding domain (DBD), oligomerization domain (OD or HR-A/B), nuclear localization signal, nuclear export signal, activator motif (AHA motif), and repressor domain (Nover et al. 2001; Scharf et al. 2012). The HR-A/B is connected to the DBD by a flexible linker of variable length. Based on the sequence characteristics of the flexible linker and HR-A/B, plant HSFs can be characterized as class A, B, or C (Chauhan et al. 2011; Guo et al. 2008; Nover et al. 2001). Because of the functional importance of HSF genes, genome-wide analyses have been conducted in different plants (Yu et al. 2022), such as *Solanum lycopersicum* (Yang et al. 2016), *Arabidopsis thaliana* (Guo et al. 2008), *Oryza sativa* (Chauhan et al. 2011; Guo et al. 2008), *Glycine max* (Li et al. 2014), *Triticum aestivum* (Ye et al. 2020), *Zea mays* (Zhang et al. 2020), *Populus trichocarpa* (Scharf et al. 2012), and *Vitis vinifera* (Hu et al. 2016). Of the 21 HSFs in *Arabidopsis*, 15 belong to Class A, 5 belong to Class B, and 1 belongs to Class C (Guo et al. 2008; Nover et al. 2001; Scharf et al. 2012).

The genome of *S. miltiorrhiza* has been sequenced (Song et al. 2020; Xu et al. 2016). However, biotic and abiotic stresses, pathogen infections, and the degeneration of elite cultivars limit the yield and quality of *S. miltiorrhiza*. Therefore, it is important to predict and characterize stress resistance genes and identify molecular mechanisms of genetic improvement. In this study, we predicted HSF genes in the whole genome using the recently reannotated high-quality reference genome of *S. miltiorrhiza* (Song et al. 2020), in order to identify genes conserved among *S. miltiorrhiza, A. thaliana*, and *O. sativa*. Phylogenetic relationships, gene structures, conserved motifs, protein domain architectures, chromosomal distributions, duplication events, selection pressures, synteny, and expression profiles were also comprehensively analyzed. This study could provide valuable information elucidating the function of HSF genes in *S. miltiorrhiza*, the adaptation mechanism of medicinal plants, and the regulatory mechanism of secondary metabolite biosynthesis. Our findings may contribute to improving the stress resistance of *S. miltiorrhiza* by informing molecular assisted breeding and intelligent breeding.

## Materials & Methods

### Sequence retrieval and preprocessing

The *S. miltiorrhiza* genome database (DSS3 v1.0) was downloaded from NGDC (2022). The current *Arabidopsis* (*A. thaliana*), rice (*O. sativa*), tomato (*S. lycopersicum*), poplar (*P. trichocarpa*), and grape (*V. vinifera*) genome databases were downloaded from Phytozome v13 (Goodstein et al. 2012). The HMM profile of the DBD domain (PF00447) was obtained from Pfam (Mistry et al. 2021). The 21 and 25 HSF protein sequences reported in *Arabidopsis* and rice were retrieved from the TAIR (Cheng et al. 2017) and RGAP (Ouyang et al. 2007) databases, respectively. The representative protein sequences in Danshen (DSS3 v1.0), *Arabidopsis* (Araport11), rice (v7.0), tomato (ITAG4.0), poplar (v4.1), and grape (v2.1) were used for the verification of putative HSF genes.

### Prediction of HSF genes in *S. miltiorrhiza* using the HBIAM method

To accurately predict the HSF genes in the *S. miltiorrhiza* genome, we created a simple, flexible, and user-friendly method called HBIAM that combines HMMER (Mistry et al. 2013), BLAST (Camacho et al. 2009), InterPro (Blum et al. 2021), alignment, and manual curation. The workflow of HBIAM follows the Input, Process, Output programming method (Fig. 1). Yellow boxes represent input data, green boxes represent output data, and blue boxes specify individual steps, with the employed programs given in brackets for each step. Following this workflow, the output data can be used as input for the subsequent BLAST search, InterPro annotation, alignment, and manual curation. The actual input data for the prediction of HSF genes in *S. miltiorrhiza* through the HBIAM method requires three input files (HMM model: PF00447, protein sequences of the *S. miltiorrhiza* genome, and the reference *Arabidopsis* HSF protein sequences), shown in red font. All of the E-values of the HMMER and the BLAST search steps are less than 10^−5^. To find more putative target genes, a blastp search is performed after the hmmsearch, using the reference *Arabidopsis* sequences. The final manually curated results are the final data used for the characterization of the HSF genes in *S. miltiorrhiza*. As the program InterProScan5 (Jones et al. 2014) runs for a relatively long time, the HBIAM method cannot be completed in one step. Therefore, starting from the verification of the InterPro annotation step, the method must be conducted step by step, and the process may need to be cycled to improve the reference *Arabidopsis* sequence. In addition, each sequence must be tested during the alignment step, and redundant sequences or sequences containing incomplete domains must be deleted during manual curation. By specifying protein sequences of different genomes and employing the HMM model for other gene families and the corresponding reference sequences, the HBIAM method can be easily applied to more species and gene families.

**Figure 1.**
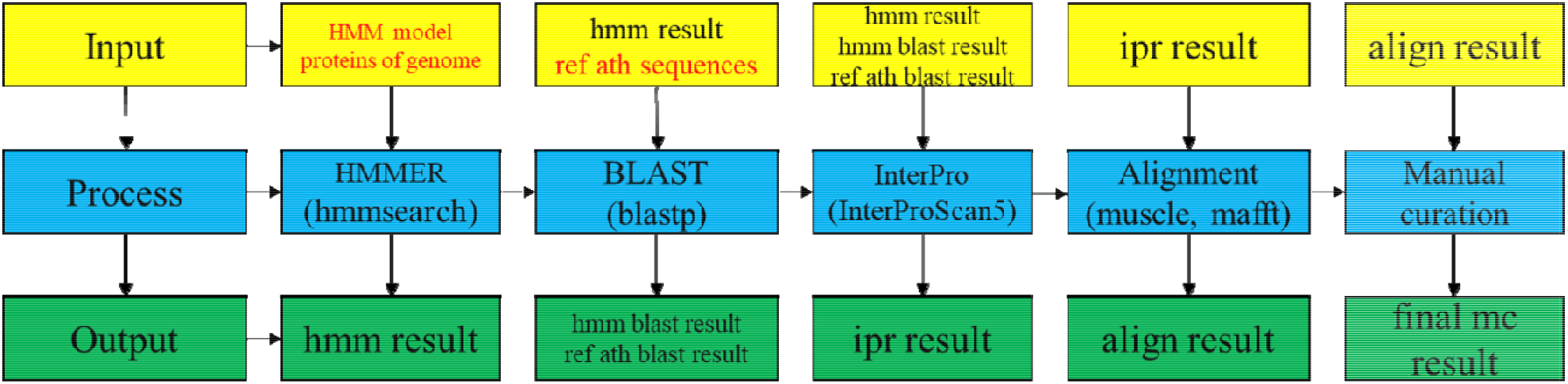
The workflow of HBIAM method.

### Sequence alignment and phylogenetic relationships of HSF genes in *S. miltiorrhiza*

To analyze the phylogenetic relationships of HSF genes in *S. miltiorrhiza*, the predicted HSF protein sequences were aligned using MUSCE v3.8.31 (Edgar 2004) with default parameters and MAFFT v7 (Katoh & Standley 2013) with the L-INS-i strategy. The neighbor-joining (NJ) tree of the HSF proteins was built using MEGA11 (Tamura et al. 2021) software, with 1000 bootstrap repetitions. The maximum-likelihood (ML) tree of the HSF proteins was built using IQ-TREE v2.20 (Minh et al. 2020) software, based on the JTT+I+G model, with 1000 bootstrap repetitions. Phylogenetic trees were annotated using the ITOL v6 Web server (Letunic & Bork 2021).

### Sequence characteristics of HSF genes in *S. miltiorrhiza*

The structures of the HSF genes were determined using the annotation file (gff3 format) of the *S. miltiorrhiza* genome. The domains of the HSF protein sequences were annotated using InterPro with InterProScan5 (Jones et al. 2014). The motifs of the HSF protein sequences were identified using Multiple EM for Motif Elicitation (Bailey et al. 2015). The cis-acting elements were detected using the plantCARE database (Lescot et al. 2002). The gene structures, protein motif structures, protein domain architectures, and cis-acting elements were all visualized using TBtools (Chen et al. 2020a).

### Chromosomal locations, gene duplications, and synteny analyses of HSF genes in *S. miltiorrhiza*

The chromosomal locations of the HSF genes were collected from the annotation file (gff3 format) of the *S. miltiorrhiza* genome (Song et al. 2020). The potential gene duplication events and synteny of the HSF genes were detected via MCScanX (Multiple Collinearity Scan toolkit) (Wang et al. 2012) software, using the default settings. The chromosomal locations in the genome and the microsyntenic relationships of the HSF genes were visualized using Circo v0.69-9 software (Krzywinski et al. 2009). To illustrate the syntenic relationships of the homologous HSF genes in *S. miltiorrhiza* and other plants, syntenic analysis maps were drawn using Dual Systeny Plot software in TBtools software (version 1.098) (Chen et al. 2020a). Synonymous (Ks) and nonsynonymous (Ka) substitution rates were calculated using WGDi (Sun et al. 2021) and PAML4 (Yang 2007), as previously described (Meng et al. 2015). For each homologous gene pair, the approximate divergence time (T; million years ago, Mya) of the duplication events was estimated using the averaged Ks values from T = Ks/2λ. The averaged Ks rate in *S. miltiorrhiza* (λ) is 6 × 10^−9^ (Li et al. 1987; Wolfe et al. 1989a; Wolfe et al. 1989b).

### Expression profiles

RNA-seq datasets (accession numbers “SRP134076” and “SRP065934”) were downloaded and selected from the NCBI SRA database and used to explore the expression patterns of HSF genes in *S. miltiorrhiza* (Wei et al. 2018; Zhang et al. 2016). SRP134076 (Wei et al. 2018) contains the RNA-seq reads of three drought-stressed and well-watered plants of *S. miltiorrhiza* obtained on day 25 (before drought) and day 31 (after six days of drought). SRP065934 (Zhang et al. 2016) contains the transcriptional profiles of eight samples of *S. miltiorrhiza* leaf callus cells subjected to salicylic acid (SA) induction for 0 h, 2 h, or 8 h. For expression analysis, the reference transcriptome of *S. miltiorrhiza* was collected from NGDC (2022). To estimate the expression profile of each sample, after passing strict quality control, the reads of these two experiments were aligned with the reference transcriptome using kallisto (Bray et al. 2016). The counts of each transcript were normalized to transcripts per million mapped reads (TPM), and the expression data were expressed as log2-based TPM values. A heatmap of the HSF gene expression patterns was drawn using the R heatmap.2 function.

## Results

### Prediction of HSF proteins in *S. miltiorrhiza*

HSF proteins, characterized by the presence of the conserved DBD and the adjacent HR-A/B, have been previously identified in the *Arabidopsis* genome (Guo et al. 2008; Nover et al. 2001). In this study, using the HSF DBD HMM model PF00447 from Pfam and the HSF protein sequences of *Arabidopsis* as input, we identified 34 putative HSF proteins in the current *S. miltiorrhiza* genome via the HBIAM method (Table 1). Each predicted HSF protein contained a typical DBD. The lengths of the predicted HSF proteins ranged from 187 aa for *SmHSF20* and *SmHSF28* to 508 aa for *SmHSF12*. One hundred and nineteen HSF genes from the five representative plant species (four dicots: *Arabidopsis* (21), tomato (26), poplar (29), and grape (18); one monocot: rice (25)) were also predicted and verified in the current genomes in Phytozome v13 (Goodstein et al. 2012) (Table S1).

**Table 1.** Putative HSF genes predicted in the Salvia miltiorrhiza genome.

| Gene Symbol | Gene locus | Group Name | Chromosome Name | Gene Start (bp) | Gene End (bp) | length (AA) |
| --- | --- | --- | --- | --- | --- | --- |
| *SmHSF01* | GWHGAOSJ001277 | A9 | GWHAOSJ00000341 | 28731 | 30471 | 293 |
| *SmHSF02* | GWHGAOSJ001304 | A7 | GWHAOSJ00000845 | 32515 | 34253 | 332 |
| *SmHSF03* | GWHGAOSJ001563 | A2 | GWHAOSJ00000135 | 89228 | 91630 | 362 |
| *SmHSF04* | GWHGAOSJ003446 | B2 | Sm01 | 52617169 | 52619131 | 290 |
| *SmHSF05* | GWHGAOSJ003593 | A9 | Sm01 | 55066677 | 55068890 | 364 |
| *SmHSF06* | GWHGAOSJ004497 | B4 | Sm01 | 66173487 | 66175033 | 341 |
| *SmHSF07* | GWHGAOSJ004649 | A7 | Sm01 | 67984109 | 67988066 | 357 |
| *SmHSF08* | GWHGAOSJ004920 | B3 | Sm01 | 70754509 | 70757262 | 233 |
| *SmHSF09* | GWHGAOSJ006555 | A2 | Sm02 | 849820 | 851921 | 316 |
| *SmHSF10* | GWHGAOSJ007410 | B2 | Sm02 | 9983519 | 9985373 | 345 |
| *SmHSF11* | GWHGAOSJ007831 | A7 | Sm02 | 17108630 | 17113725 | 334 |
| *SmHSF12* | GWHGAOSJ008020 | A1 | Sm02 | 20136211 | 20139358 | 508 |
| *SmHSF13* | GWHGAOSJ010875 | A6 | Sm03 | 12486179 | 12487783 | 328 |
| *SmHSF14* | GWHGAOSJ011087 | A5 | Sm03 | 15038175 | 15041500 | 482 |
| *SmHSF15* | GWHGAOSJ012410 | A3 | Sm03 | 36193660 | 36195130 | 352 |
| *SmHSF16* | GWHGAOSJ014590 | A4 | Sm04 | 38679959 | 38681485 | 388 |
| *SmHSF17* | GWHGAOSJ014694 | A9 | Sm04 | 41053586 | 41054446 | 265 |
| *SmHSF18* | GWHGAOSJ014718 | A9 | Sm04 | 41452448 | 41453274 | 252 |
| *SmHSF19* | GWHGAOSJ014732 | A9 | Sm04 | 41746782 | 41747660 | 260 |
| *SmHSF20* | GWHGAOSJ014740 | B5 | Sm04 | 41858053 | 41862419 | 187 |
| *SmHSF21* | GWHGAOSJ015793 | B4 | Sm04 | 59854187 | 59855666 | 290 |
| *SmHSF22* | GWHGAOSJ017002 | A4 | Sm04 | 73026010 | 73028332 | 425 |
| *SmHSF23* | GWHGAOSJ017966 | A1 | Sm05 | 15867302 | 15872359 | 502 |
| *SmHSF24* | GWHGAOSJ018248 | A8 | Sm05 | 32251902 | 32254411 | 383 |
| *SmHSF25* | GWHGAOSJ018838 | B2 | Sm05 | 44095866 | 44097284 | 227 |
| *SmHSF26* | GWHGAOSJ020379 | A6 | Sm05 | 62362106 | 62363895 | 313 |
| *SmHSF27* | GWHGAOSJ021001 | A4 | Sm06 | 4636102 | 4638552 | 399 |
| *SmHSF28* | GWHGAOSJ021164 | B5 | Sm06 | 6211345 | 6212895 | 187 |
| *SmHSF29* | GWHGAOSJ021890 | C1 | Sm06 | 15562431 | 15563668 | 336 |
| *SmHSF30* | GWHGAOSJ023505 | C1 | Sm06 | 59448677 | 59452050 | 353 |
| *SmHSF31* | GWHGAOSJ026491 | A3 | Sm07 | 57125201 | 57127729 | 487 |
| *SmHSF32* | GWHGAOSJ027753 | A9 | Sm08 | 8009395 | 8011247 | 341 |
| *SmHSF33* | GWHGAOSJ028128 | B3 | Sm08 | 14535073 | 14537761 | 224 |
| *SmHSF34* | GWHGAOSJ028526 | B1 | Sm08 | 28546765 | 28550994 | 269 |

### Phylogenetic analysis of HSF genes in *S. miltiorrhiza*

To characterize the evolutionary relationships of the HSF genes in *S. miltiorrhiza*, NJ and ML phylogenetic trees were constructed, based on the full-length protein sequence alignment of the HSF genes in *S. miltiorrhiza, A. thaliana* and *O. sativa*. Based on prior evolutionary analyses and the classification criteria previously used to characterize *A. thaliana* and other plants (Chauhan et al. 2011; Guo et al. 2008; Nover et al. 2001), the HSF genes in *S. miltiorrhiza* were divided into three classes: Class A (22), Class B (10), and Class C (2) (Fig. 2). Class A was further divided into nine groups (Groups A1, A2, A3, A4, A5, A6, A7, A8, and A9), which contained 2, 2, 2, 3, 1, 2, 3, 1, and 6 HSF genes, respectively. Class B was divided into four groups (Groups B1, B2, B3, B4, and B5), which contained 1, 3, 2, 2, and 2 HSF genes, respectively. Class C contained Group C1, which included only 2 HSF genes. Class A contained the most HSF genes, while Class C contained the fewest.

**Figure 2.**
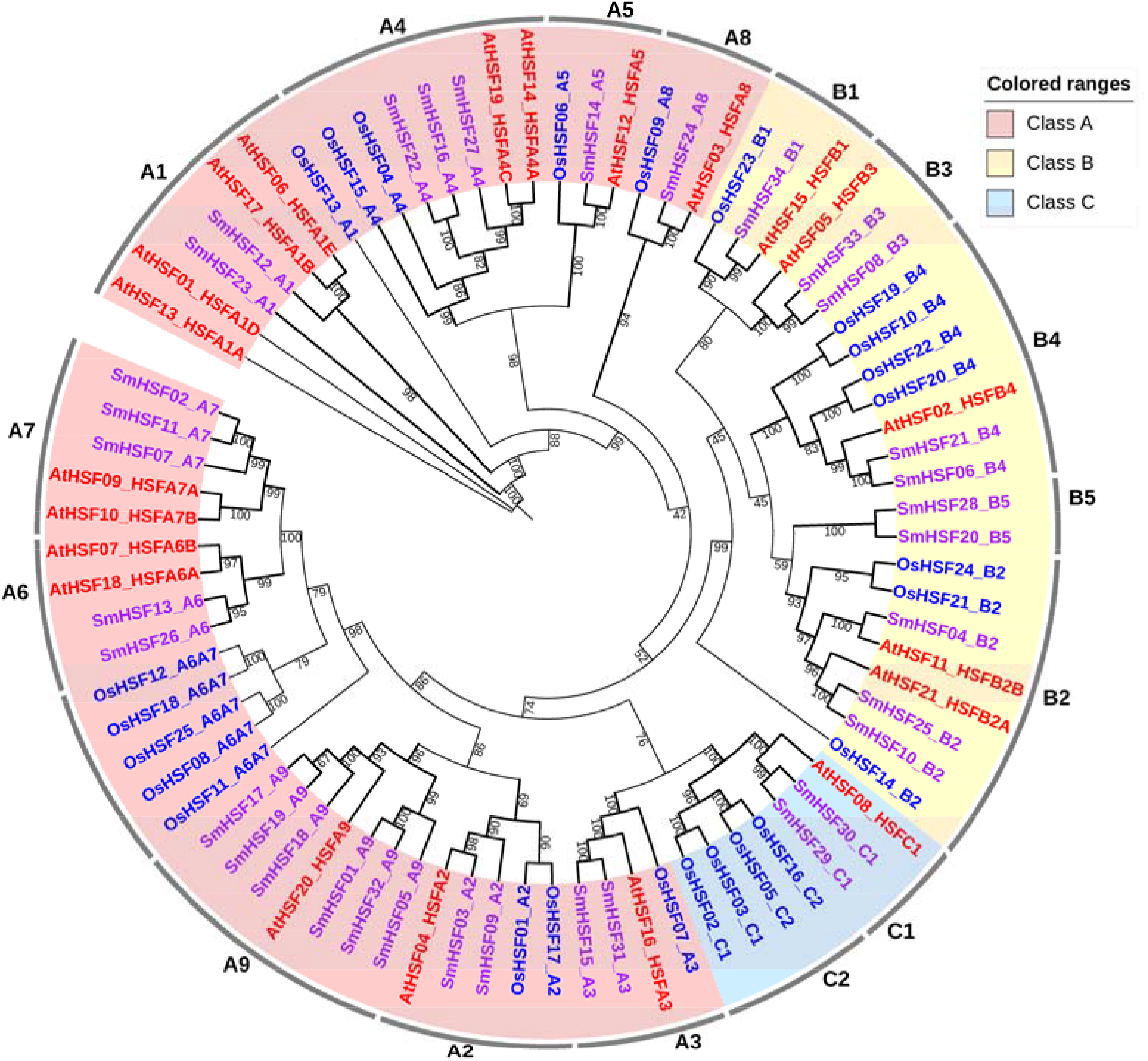
The evolutionary relationships of the HSF genes in S. miltiorrhiza, A. thaliana and O. sativa. The three HSF gene classes (A, B, and C) were shown by three underground colors. The font colors of HSF ID from S. miltiorrhiza, A. thaliana and O. sativa were purple, red and blue respectively.

### Sequence characteristics of HSF genes in *S. miltiorrhiza*

To investigate the structural diversity of the HSF genes in *S. miltiorrhiza*, the exon-intron structures of the HSF genomic sequences, domain architectures, and conserved motifs of the HSF proteins were compared according to the phylogenetic relationships. Almost all of the HSF genes in *S. miltiorrhiza* contained fewer exons, usually two exons, one short and the other long. *SmHSF11* and *SmHSF07* in Group A7 had a third, shorter exon (Fig. 3A). Closely related HSF genes in the same group had similar exon-intron structures, and those with closer evolutionary relationships exhibited greater similarity in the number and length of exons and introns. Domains other than the DBD were rarely found in the C-terminal regions (Fig. 3B). Twenty-four conserved motifs (motif1-motif24) were predicted; these motifs were specific to each group (Fig. 3C). The composition of the structural motifs varied among the different HSF classes or groups, while the motifs within each group were similar. Additionally, the motifs encoding the DBD in the N-terminal regions were relatively conserved, suggesting that the functions of the HSF proteins may be intergroup-specific and intragroup-conserved. The HSF genes in *S. miltiorrhiza* from the same group generally had similar structural and sequence characteristics (Fig. 3). Using plantCARE, 11 cis-elements were detected, and a 2 kb region upstream of the translation start site was selected (Fig. 4). These cis-elements may be involved in the regulation of hormone responsiveness and environmental stress.

**Figure 3.**
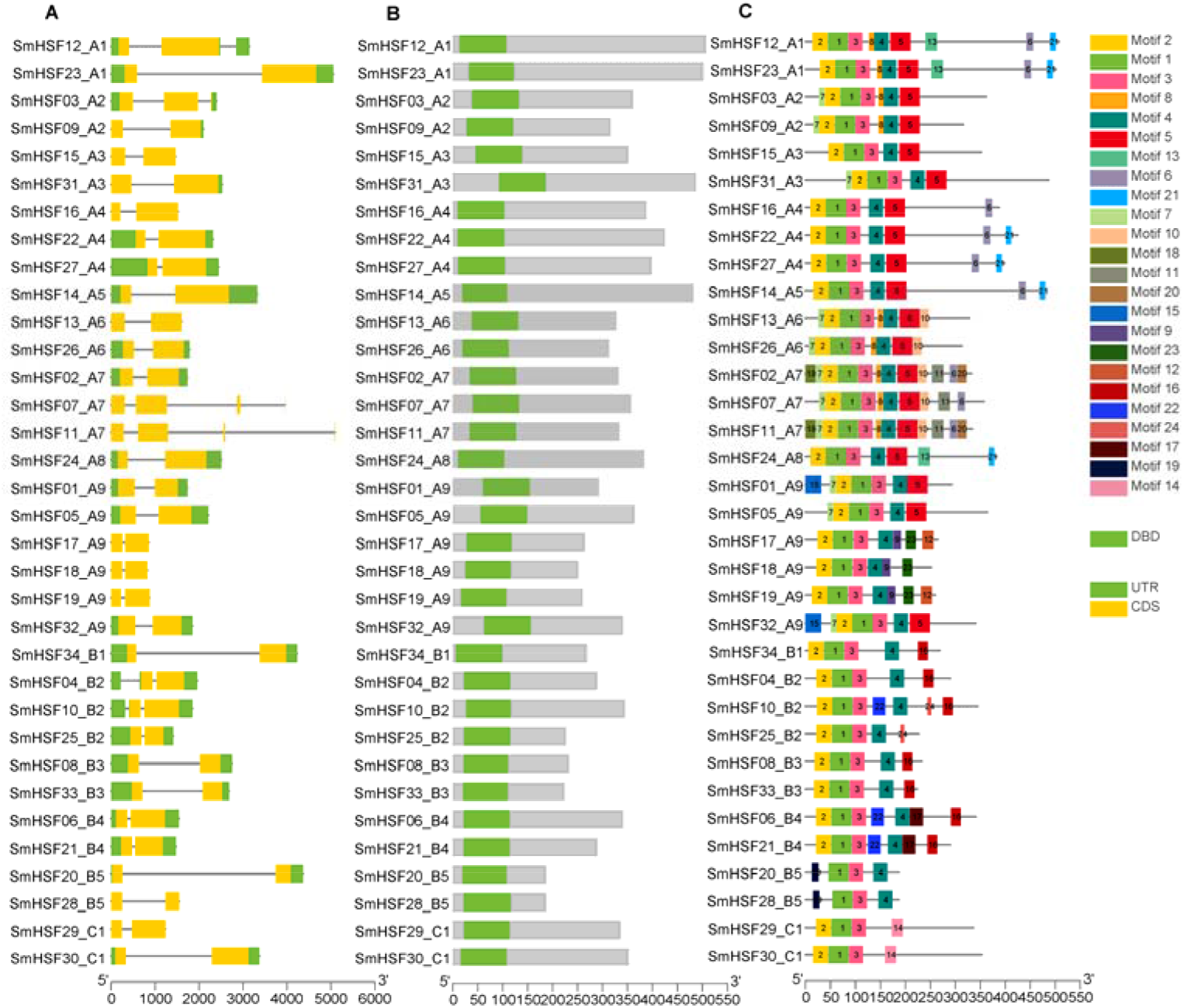
Sequence characteristics of HSF genes in S. miltiorrhiza. (A) The exon-intron structures of HSF genes in S. miltiorrhiza. (B) The protein domain architectures of HSF genes in S. miltiorrhiza. (C) The conserved motifs of the HSF proteins in S. miltiorrhiza.

**Figure 4.**
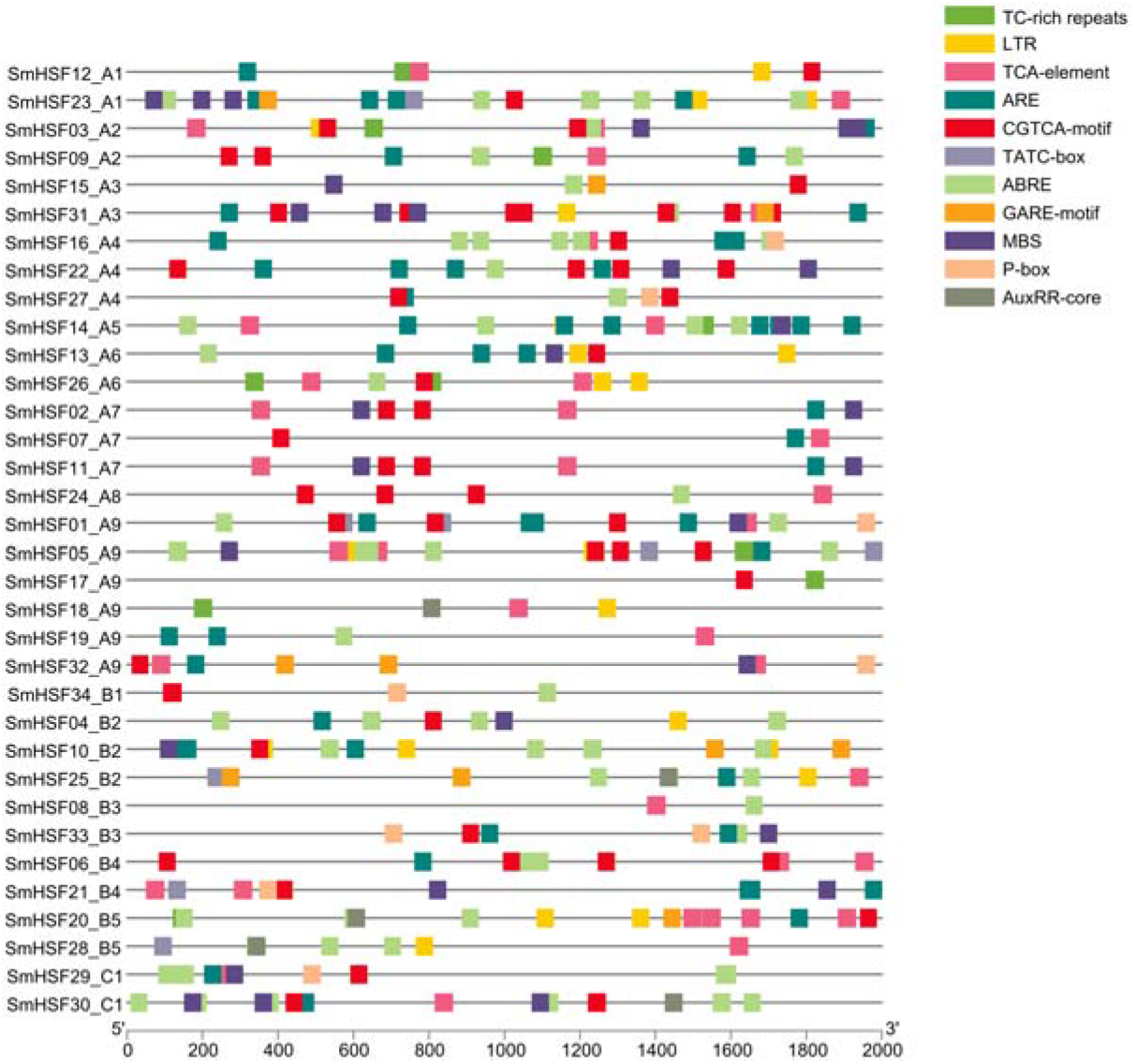
The Cis-acting elements of HSF genes in S. miltiorrhiza.

### Chromosome location and gene duplication of HSF genes in *S. miltiorrhiza*

To investigate the chromosome distributions of the predicted HSF genes in *S. miltiorrhiza*, the chromosomal locations of the 34 HSF genes were extracted from the annotation of *S. miltiorrhiza* under strict quality control. Most of the HSF genes were distributed across eight chromosomes at different densities: Sm1 to Sm8 comprised 5, 4, 3, 7, 4, 4, 1, and 3 HSF genes, respectively (mean: 4; max: Sm4, 7; min: Sm7, 1) (Fig. 5, Table 1). Only three HSF genes were not specified. To examine the gene duplication events, all the genes were analyzed using BlastP (Camacho et al. 2009) and MCScanX software (Wang et al. 2012). Thirteen segmental duplication events were confirmed in the whole genome of *S. miltiorrhiza* (Fig. 5, Table S2). To examine selection pressure, the Ks, Ka, and Ka/Ks ratio of the HSF gene pairs were calculated (Table S2). The Ka/Ks ratio was estimated to be less than 1, indicating that the duplicated HSF genes in *S. miltiorrhiza* were under strong negative selection. Gene duplication analysis indicated that the segmental duplication events contributed to the expansion of the *S. miltiorrhiza* HSF gene family.

**Figure 5.**
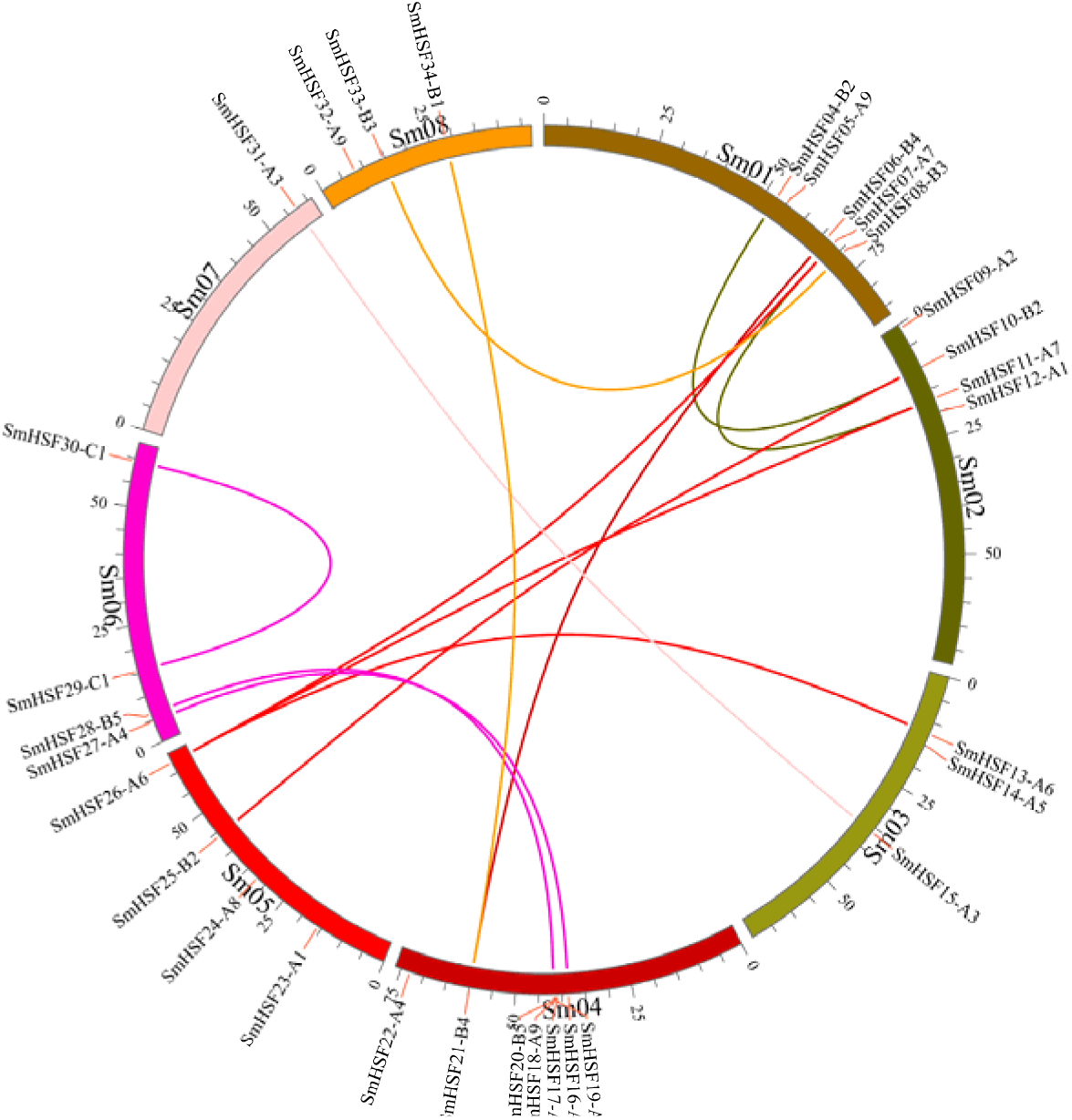
Chromosomal locations of HSF genes and duplicated gene pairs in the S. miltiorrhiza genome. Chromosomes 1-8 are shown with different colors and in a circular form. The approximate distribution of each HSF genes in the S. miltiorrhiza genome is marked on the circle with a short red line. Colored curves denote the details of syntenic regions between HSF genes in the S. miltiorrhiza genome.

To investigate the phylogenetic mechanisms of the *S. miltiorrhiza* HSF genes, a comparative syntenic analysis (Fig. 6) of *S. miltiorrhiza* and the five other representative plants, including four dicots (*P. trichocarpa, S. lycopersicum, A. thaliana*, and *V. vinifera*) and one monocot (*O. sativa*), was performed. A total of 24 (29, 82.76%), 20 (26,76.92%), 17 (21, 80.95%), 16 (18, 88.89%), and 10 (25, 40.00%) HSF genes in *P. trichocarpa, S. lycopersicum, A. thaliana, V. vinifera*, and *O. sativa*, respectively, were involved in the formation of 39, 35, 23, 23, and 12 orthologous HSF gene pairs, respectively (Table S3).

**Figure 6.**
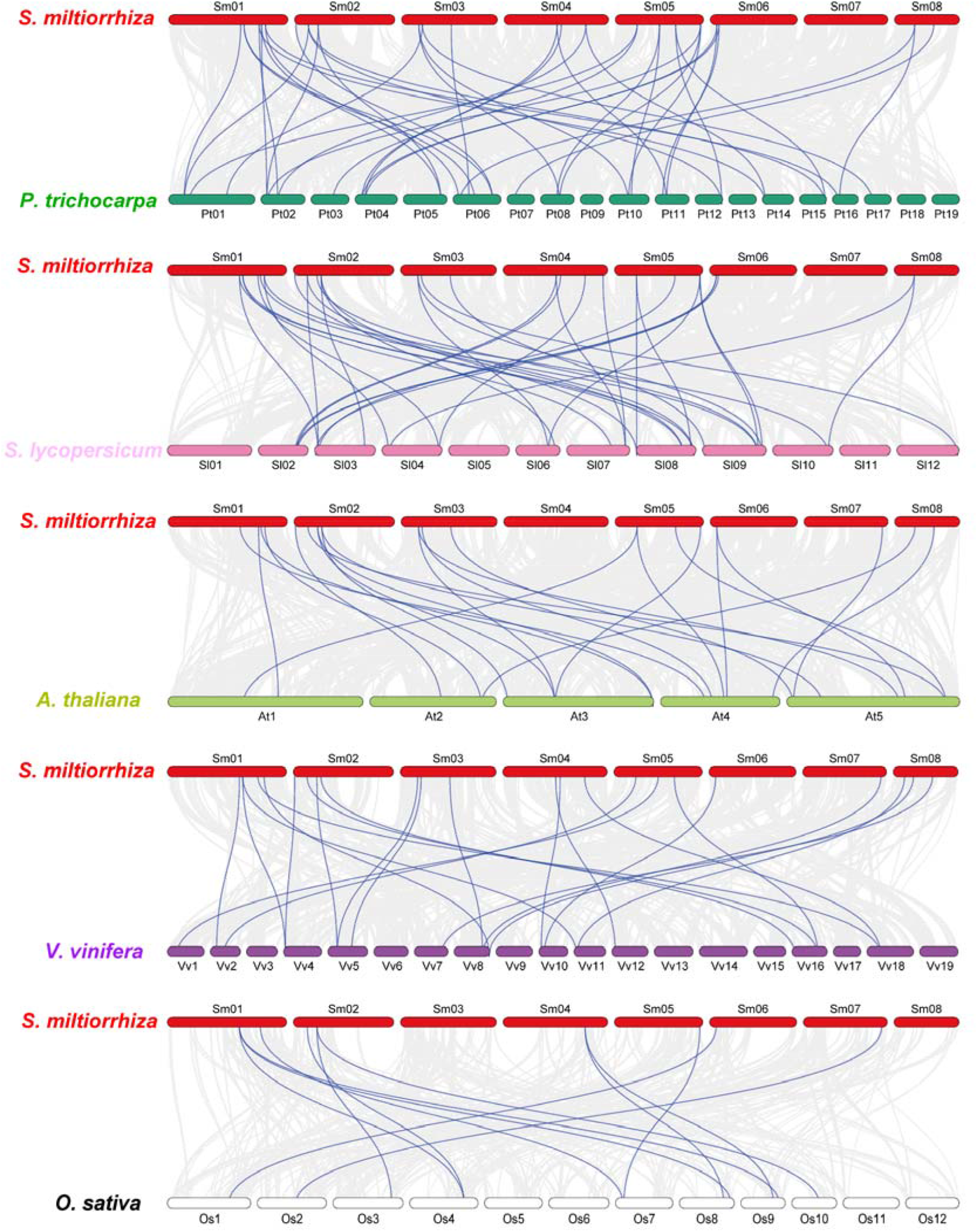
Synteny analysis of HSF genes in S. miltiorrhiza and five representative plant species. Gray lines in the background denote collinear blocks within S. miltiorrhiza and other plant genomes, and blue lines highlight the syntenic HSF gene pairs, with five representative plant species being *P. trichocarpa, S. lycopersicum, A. thaliana, V. vinifera*, and *O. sativa*.

### Expression of *S. miltiorrhiza* HSF genes under drought stress and SA induction

To explore the potential functions of *S. miltiorrhiza* HSF genes, we used public RNA-seq data from two experiments related to drought stress (Wei et al. 2018) and SA induction (Zhang et al. 2016). The 34 *S. miltiorrhiza* HSF genes were detected in all samples at the gene level (Fig. 7, Table S4 and Table S5). These HSF genes displayed differential expression in response to drought stress or SA induction. Under drought stress, most HSF genes in groups A1, A4, A5, B1, B2, B3, and C1 were highly expressed, such as *SmHSF22, SmHSF10*, and *SmHSF30. SmHSF05* and *SmHSF032* in group A9 were also highly expressed (Fig. 7A and Table S4). Groups A1, A2, A3, A4, A5, A6, A7, A8, A9, and C1 contained at least one HSF gene that was highly expressed in response to SA induction, while almost all HSF genes in groups B1, B2, B3, and B5 were highly expressed (Fig. 7B and Table S5). Most of the HSF genes in plants subjected to SA induction for 2 h showed a high expression level, a clearly different expression pattern when compared with the similar expression pattern of the CK and SA-8h, such as *SmHSF27* in group A4 (Fig. 7B).

**Figure 7.**
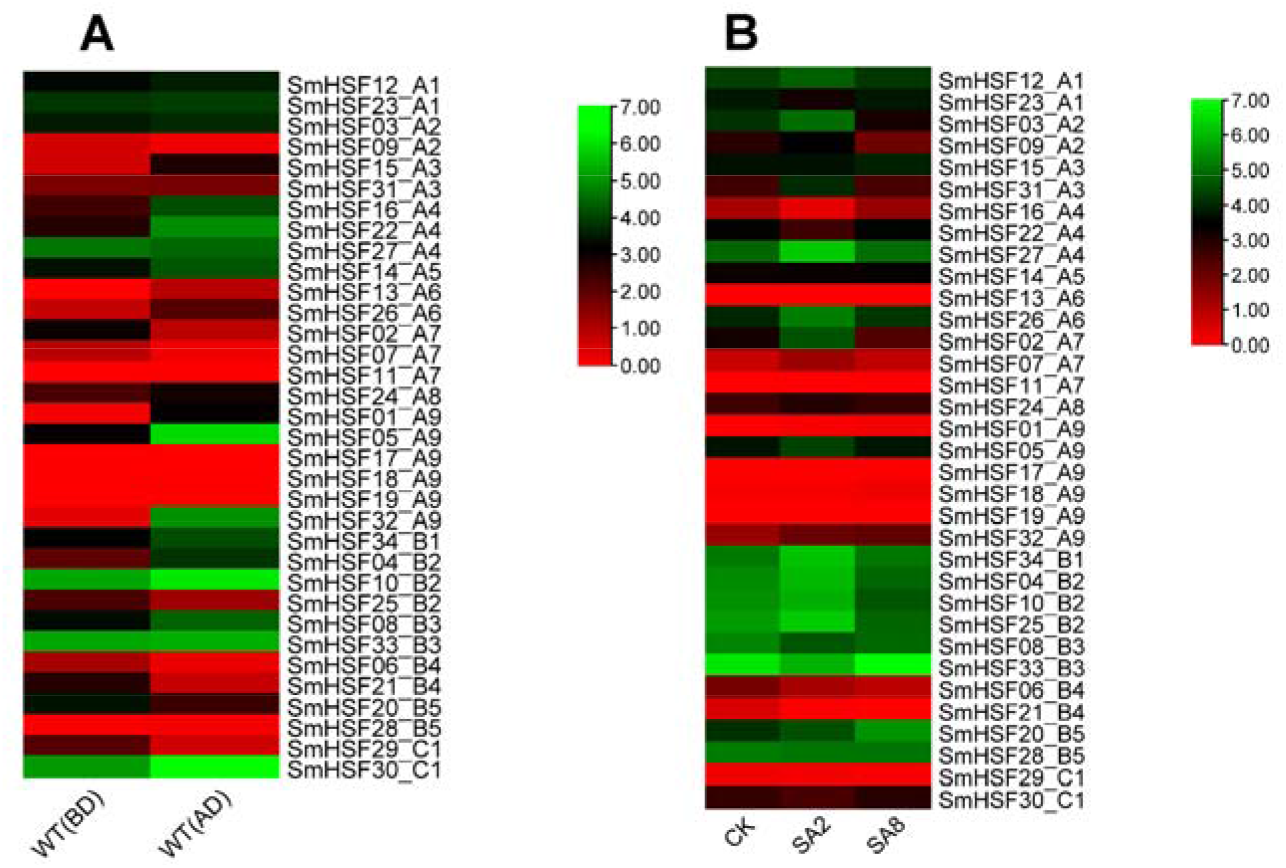
Heatmap of HSF gene expression patterns under drought stress and SA induction.

## Discussion

In *A. thaliana*, HSF genes play important roles in diverse plant growth (Gonzalez-Bayon et al. 2019), developmental processes, (Fortunati et al. 2008) and physiological processes (Andrasi et al. 2021; Faragó et al. 2018). The complex functional characteristics of HSF genes facilitate their responses to heat (Cortijo et al. 2017), drought (Zhu et al. 2020), cold (Olate et al. 2018), salt (Bian et al. 2020; Jing et al. 2019), anoxia (Banti et al. 2010), heavy metal (Chen et al. 2020b; Shim et al. 2009), irradiation (Yamanouchi et al. 2002), and pathogens (Zhou et al. 2018). In this study, we precisely predicted the HSF genes in the *S. miltiorrhiza* genome to identify genes conserved in *S. miltiorrhiza, A. thaliana*, and *O. sativa*.

We identified 34 putative HSF genes in the *S. miltiorrhiza* genome. The number of HSF genes in *S. miltiorrhiza* was higher than that reported for *A. thaliana* (21) (Guo et al. 2008; Nover et al. 2001), *O. sativa* (25) (Chauhan et al. 2011; Guo et al. 2008), *S. lycopersicum* (24) (Li et al. 2014), *P. trichocarpa* (27) (Scharf et al. 2012), *V. vinifera* (Hu et al. 2016), and *Z. mays* (30) (Zhang et al. 2020), and lower than that reported for *G. max* (52) (Li et al. 2014) and *T. aestivum* (56) (Ye et al. 2020). Genome duplication events may have contributed to the increased number of HSF genes in polyploid plants, such as paleotetraploid soybean or allohexaploid wheat.

In the 34 putative HSF proteins identified in *S. miltiorrhiza*, the general sequence characteristics, including gene structure, conserved motifs, and protein domain architectures, were similar to those observed in previous studies. Two group B5 genes (*SmHSF20* and *SmHSF28*) and two group C1 genes (*SmHSF29* and *SmHSF30*) present in *A. thaliana* and *O. sativa* were also present in *S. miltiorrhiza*, while group C2 genes were not present. Each class A group contained at least one HSF gene in *S. miltiorrhiza*. Moreover, the ML phylogenetic tree illustrated that groups A4 and A5 belonged to the same clade, and groups A6 and A7 belonged to the same clade. Therefore, we concluded that there were 13 phylogenetic groups of HSF genes in these plants (groups A1-A3, group A4A5, group A6A7, group A8, group A9, groups B1-B5, and group C1C2). The paralogous HSF genes in a phylogenetic group often possess identical or similar gene structures and domain organizations and might bind or interact with identical or similar target proteins. The different phylogenetic groups of HSF proteins might contribute to the development of various HSF gene functions related to growth, development, and responses to biotic and abiotic stresses.

Segmental duplication is an important evolutionary mechanism that accelerates the rapid expansion of gene families. This mechanism has been detected in most plant genomes, as most plant species are diploidized polyploids, and numerous duplicated chromosomal blocks are present in the current genomes (Panchy et al. 2016; Qiao et al. 2019; Zhu et al. 2014). Thirteen segmental duplication events, involving 67.65% of the HSF genes (23/34), were confirmed in the whole genome of *S. miltiorrhiza* (Table S2), indicating that segmental duplication contributed to the expansion of the *S. miltiorrhiza* HSF gene family. To evaluate selection pressure, the Ks, Ka, and Ka/Ks ratio of the HSF gene pairs were calculated (Table S2). The Ka/Ks ratio was estimated to be less than 1, indicating that the duplicated HSF genes in *S. miltiorrhiza* were under negative selection pressure. By calculating the divergence time of the duplicated HSF gene pairs in the *S. miltiorrhiza* genome, we inferred that seven duplicated pairs of paralogous HSF genes were derived from a recent duplication event, and four duplicated pairs of paralogous HSF genes were derived from an ancient duplication event during the shaping of the *S. miltiorrhiza* genome (Table S2). Thus, segmental duplications were the primary driving forces behind the evolution of HSF genes during plant speciation and gene functionalization.

To further understand the potential functions of HSF genes under drought stress (Wei et al. 2018) and SA induction (Zhang et al. 2016) in *S. miltiorrhiza*, we conducted an expression pattern analysis of publicly available RNA-seq data. Most of the HSF genes displayed differential expression (Figure 7). The expression of most HSF genes increased after drought stress was experienced (Figure 7A), indicating that these HSF genes may participate in the drought resistance of *S. miltiorrhiza*. In plants subjected to SA induction for 2 h, the expression level of most HSF genes increased, while that of several HSF genes decreased. The expression of almost all genes returned to a normal level when the plants were subjected to SA induction for 8 h (Fig. 7B). This differential expression pattern suggested that the HSF genes were involved in SA-mediated adaptive stress resistance. Further investigations verifying the functions of HSF genes in *S. miltiorrhiza* would provide more insights into the regulatory mechanisms of these genes in growth, development, and stress responses.

## Conclusions

In this study, 34 HSF genes were predicted in *S. miltiorrhiza*, and their sequence, structure, evolution, and expression characteristics were analyzed. These HSF genes were unevenly distributed across eight chromosomes. Thirteen inferred segmental duplication events involving 23 HSF genes indicated that segmental duplication contributed heavily to the expansion of the HSF gene family. Expression analysis of RNA-seq data suggested that these HSF genes may be involved in the stress resistance of *S. miltiorrhiza*. This comprehensive analysis will inform future studies of the functions of HSF genes *S. miltiorrhiza*.

