## Supplementary Information for "Gene prediction and evolutionary characterization of heat shock transcription factors in *Salvia miltiorrhiza*"

**Table S1.** 153 HSF genes predicted in *S. miltiorrhiza* and five representative plant species.

**Table S2.** Estimates of the dates for the segmental duplication events in the HSF gene pairs in *S. miltiorrhiza*.

**Table S3.** Orthologous relationships of HSF genes between *S. miltiorrhiza* and other five plant species.

**Table S4.** Log2(TPM) transcription count data for 34 HSF genes under drought stress.

**Table S5.** Log2(TPM) transcription count data for 34 HSF genes under SA induction.

**File S1.** FASTA format of the 153 full-length HSF protein sequences in *S. miltiorrhiza* and five other representative plant.
